# Human prostaglandin reductases dearomatize and inactivate benzothiazinone antitubercular drugs

**DOI:** 10.64898/2026.02.16.706187

**Authors:** Myriam Briki, Caroline Wagner, Vincent Desfontaine, Anthony Vocat, Onya Opota, Sarah Abou-Zite, Océane Corpataux, Benedetta Cena, Sylvie Guinchard, François Versace, Amarande Murisier, Thomas Mercier, Emilyne Blattes, Carine Bardinet, Paul Thoueille, Vladimir Wsol, Haithem Chtioui, Laura Rothuizen, Aimable Nahimana, Axel Bellotti, Laurent R. Chiarelli, Dany Spaggiari, Neeraj Dhar, Richa Mishra, Raphaël Sommer, Andréanne Lupien, Michel A. Duchosal, Benoît Guery, François Girardin, Giuseppina Milano, Olga Riabova, Vadim Makarov, Thierry Buclin, Stewart T. Cole, Eva Choong, Laurent A. Decosterd

## Abstract

Macozinone (MCZ, PBTZ169) is a potent clinical stage benzothiazinone antitubercular agent that covalently inhibits the essential mycobacterial flavoenzyme DprE1. In some mammals, MCZ undergoes reductive dearomatization to H_2_MCZ, a Hydride Meisenheimer Complex, identified as the major circulating metabolite in humans. We demonstrate for the first time that the NADPH-dependent human prostaglandin reductases PTGR1 and PTGR2 catalyze MCZ dearomatization into H_2_MCZ, resulting in loss of antimycobacterial activity. This reaction represents a heretofore undescribed host-mediated metabolic inactivation pathway for a therapeutic agent. Although H_2_MCZ may constitute a transient reactive intermediate, *ex vivo* and cellular data indicate that it does not contribute to DprE1 inhibition *in vivo*.

Pharmacological inhibition of PTGR1 and PTGR2 using diclofenac, indomethacin, dicumarol, or the selective inhibitor PTGR2-IN-1 suppresses H_2_MCZ formation and partially restores MCZ antimycobacterial activity *in vitro*. Together, our findings uncover a previously unrecognized noncanonical enzymatic mechanism of drug metabolism involving dearomatization in humans. Targeting prostaglandin reductases may represent a strategy to enhance benzothiazinone exposure and efficacy.

## Introduction

Tuberculosis (TB) remains one of the world’s deadliest diseases, causing 1.25 million deaths in 2023, making it the leading cause of death from a single infectious agent (surpassing COVID-19)(2024). Even more alarming is the growing number of multidrug-resistant TB (MDR-TB) cases, defined as resistant to rifampicin and isoniazid. Among the emerging treatments against both susceptible- and MDR-TB, benzothiazinones constitute an attractive class, showing potent antimycobacterial activity by covalently binding to the critical flavoenzyme decaprenylphosphoryl-β-D-ribose 2-epimerase (DprE1). The resultant blockade of arabinan polymer synthesis disrupts the mycobacterial cell wall (Makarov *et al*, 2009; Neres *et al*, 2012; Trefzer *et al*, 2012) and induces cell lysis.

Macozinone (MCZ), also known as PBTZ169 (**Fig. 1A**), is a promising, highly potent benzothiazinone derivative developed after chemical optimization from the lead compound BTZ-043 (Makarov *et al*, 2014). It has proven active against susceptible and drug-resistant *Mycobacterium tuberculosis* strains with minimum inhibitory concentration (MIC) values below 0.0005 μg/mL (Lupien *et al*, 2018). Its effectiveness has been confirmed *in vivo,* notably in guinea pig and mouse models (Makarov *et al*., 2014).

**Figure 1.**
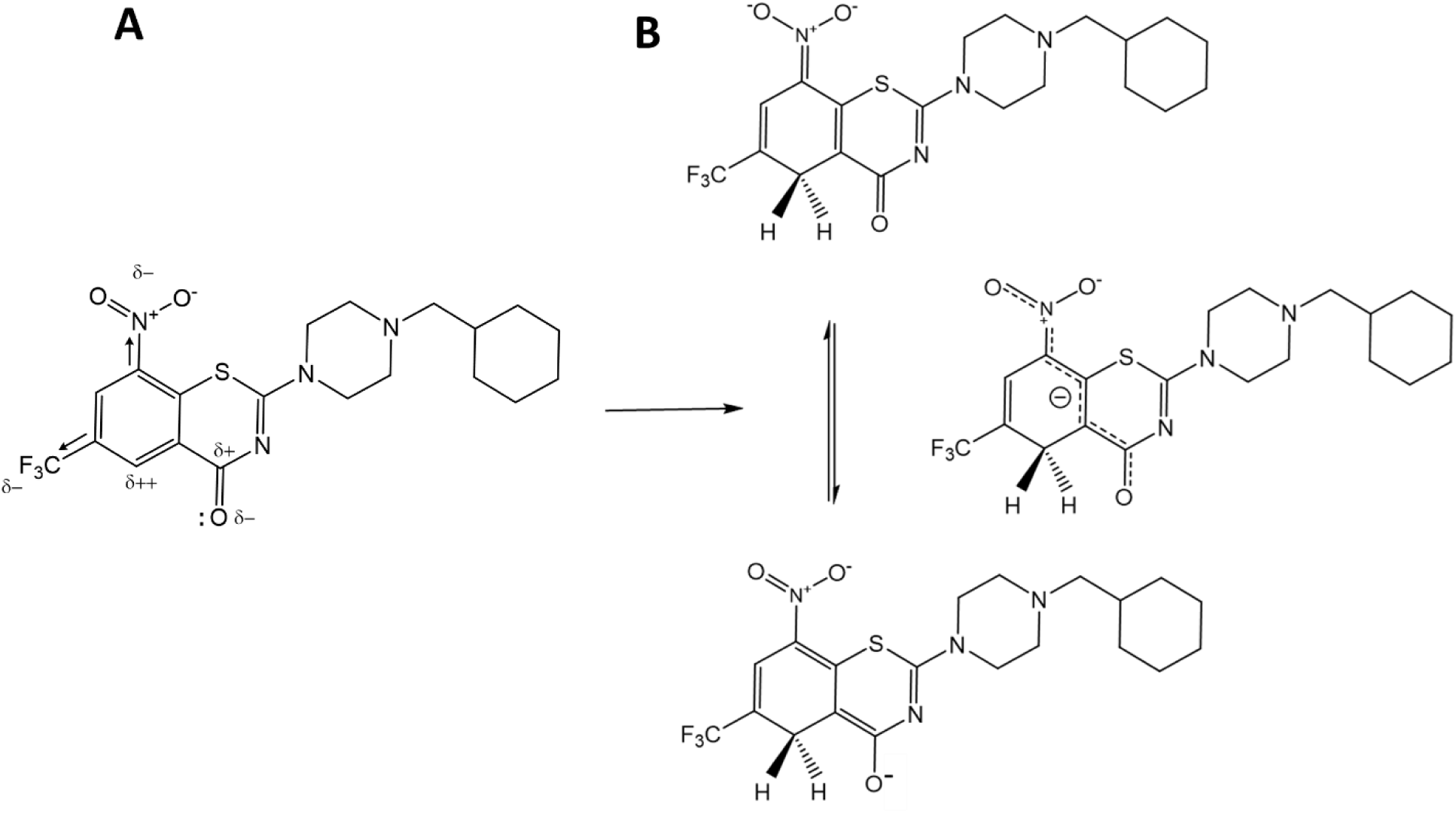
Chemical structures of (**A**) Macozinone (MCZ; PBTZ169) showing the relevant electrochemical features of the molecule. (**B**) the reduced, dearomatized, metabolite p-H_2_MCZ depicted as the two mesomeric forms in equilibrium.

Both MCZ and BTZ-043 are reductively activated to form a nitroso moiety that reacts with a key cysteine (Cys387) located in the active site of DprE1. This generates a semi-mercaptal that results in covalent inhibition(Neres *et al*., 2012).

MCZ has successfully passed preclinical safety, toxicology, pharmacokinetic as well as antimycobacterial activity assessments in Phase I and II clinical trials that took place both in Russia and in Switzerland (https://clinicaltrials.gov/study/NCT03423030 ; https://clinicaltrials.gov/study/NCT03776500). In parallel, BTZ-043, following tissue distribution studies in murine models, (Rompp *et al*, 2025) has reached Phase II trials, which demonstrated an excellent safety profile and a bactericidal activity in the range of rifampicin at a dose of 10 mg/kg (Heinrich *et al*, 2025).

In a seminal publication (Kloss *et al*, 2017), Kloss *et al*. reported that benzothiazinones (both MCZ and BTZ-043) follow an unprecedented metabolic pathway yielding a Hydride Meisenheimer Complex (HMC), which represents the first example of HMC formation in humans, and more generally in mammals. Specifically, MCZ undergoes bioreductive dearomatization *in vivo,* producing the peculiar metabolite *para*-H_2_MCZ (*p-*H_2_MCZ) (**Fig. 1B**, depicted as the two mesomeric forms), which may revert to the parent compound in the presence of oxygen (Kloss *et al*., 2017).

The regioselective reduction to *p*-H_2_MCZ indicates that the reaction is carried out enzymatically (Kloss *et al*., 2017): whereas chemical reduction by NaBH_4_ yields a mixture of *ortho*- and *para-*H_2_MCZ (*o-* and *p-*H_2_MCZ, respectively), the latter being the sole species observed *in vivo*. Explanations for this unprecedented enzymatic reaction, and its consequences for antimycobacterial activity, are lacking despite H_2_MCZ, which accumulates after multiple doses (Desfontaine *et al*, 2023), being the most abundant metabolite circulating in rats and humans (Spaggiari *et al*, 2019).

The FADH_2_/FAD redox chemistry in the DprE1 active site mediates reduction of the nitro group of benzothiazinone drugs into the reactive nitroso moiety. A potential role for the intermediate HMC metabolite has been suggested in this reaction(Liu *et al*, 2021), considering that H_2_MCZ need only lose a water molecule to generate the corresponding, putatively active nitroso species (Liu *et al*, 2021). Although disappointing to date, the search continues for benzothiazinones with a reduced HMC formation propensity that retain potent antimycobacterial activity (Joch *et al*, 2024; Keiff *et al*, 2024).

The aims of the present study were i) to identify the enzyme(s) responsible for the *in vivo* dearomatization of MCZ into the HMC H_2_MCZ; ii) to study the consequence of this metabolism on antimycobacterial activity; and iii) to assess the impact of modulation of MCZ metabolism on its antimycobacterial activity.

We successively investigated various enzymatic systems likely to catalyze MCZ dearomatization, including carbonyl-reducing enzymes (NAD(P)H:quinone oxidoreductases, and reductases from the short-chain dehydrogenase/reductase and aldo-ketoreductase superfamilies) and thiol-based antioxidant enzymes (glutathione reductase, thioredoxin reductase, and glutathione transferase). We further extended our exploration to general enzymatic pathways mediating carbon-carbon double bond reduction (Huang *et al*, 2014) in microorganisms and mammals.

## Results

### Identification of the putative macozinone reductase(s)

Our investigations were guided by results from *ex vivo* incubation experiments in mammalian tissues with prototypical inhibitors of various known reductive enzymatic pathways (**Supplementary Table S1**). Monitoring the rate of MCZ reduction to H_2_MCZ by mass spectrometry allowed us to determine the decrease in H_2_MCZ formation in the presence of inhibitors, as compared to controls. Any inhibition of the reductase systems under study opened the way to subsequent MCZ incubation experiments, ultimately with the respective pure recombinant human enzymes.

### Enzyme candidate screening

To establish the screening assay, we first tested various biological systems for their ability to reduce MCZ into H_2_MCZ *in vitro* by simultaneously monitoring their presence by high-performance liquid chromatography coupled to tandem mass spectrometry (LC-MS/MS), using assays developed (Spaggiari *et al*., 2019) and further optimized (Desfontaine *et al*., 2023) in our laboratory. Typical chromatographic profiles of H_2_MCZ in various *in vivo*, *in vitro* and *ex vivo* samples are shown in **Fig. 2A**.

**Figure 2.**
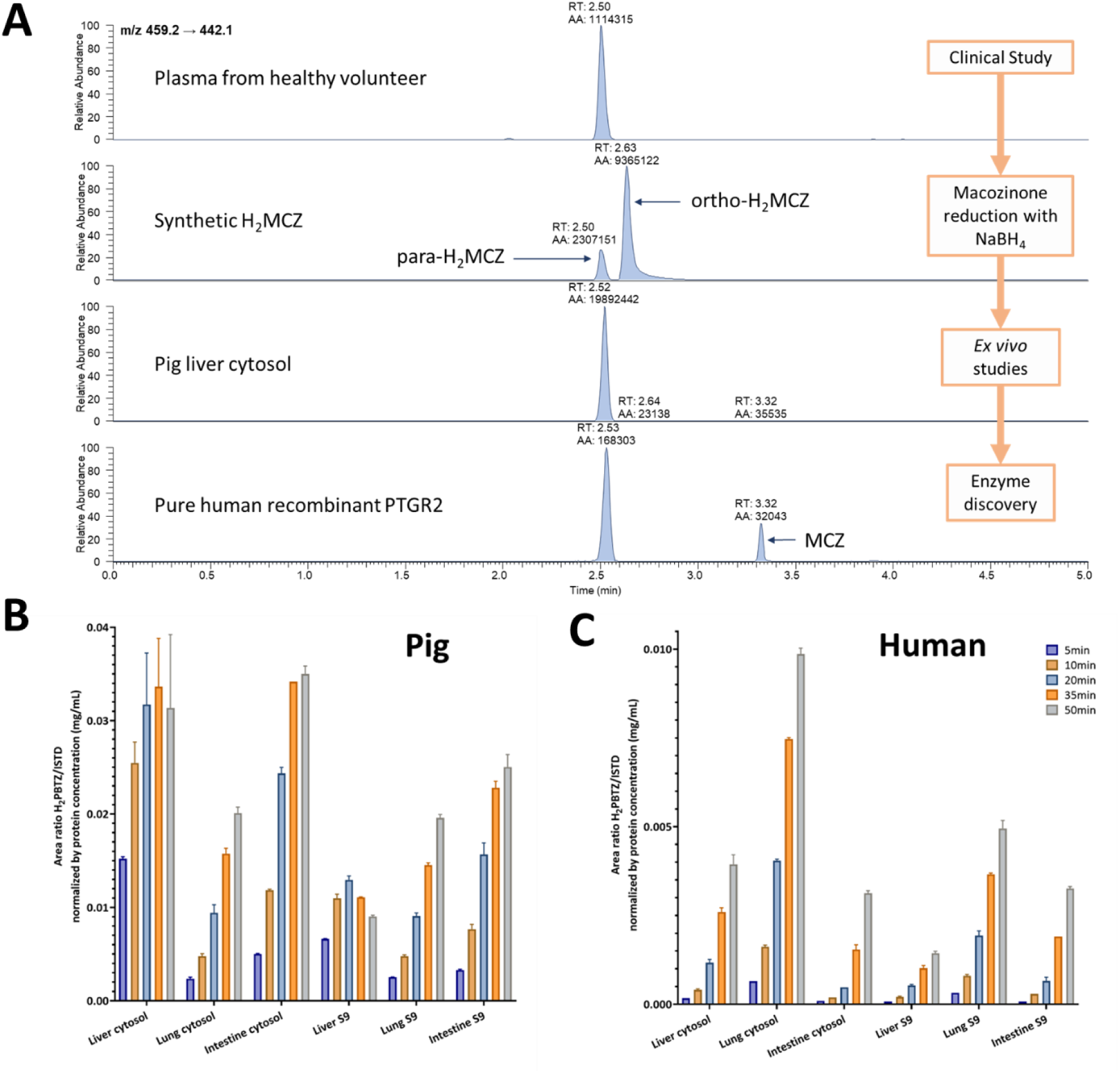
Biological systems that can reduce MCZ. (**A**) Chromatographic profiles of H_2_MCZ in various *in vitro*, *ex vivo*, and *in vivo* samples. *Ex vivo* incubation: H_2_MCZ production in (**B**) pig and (**C**) human cytosolic and S9 fractions. No H_2_MCZ was formed in microsomes nor in mitochondria from tissues. Pig spleen cytosol and S9 fractions also yielded H_2_MCZ (not shown).

Incubation of MCZ in human whole blood, peripheral blood mononuclear cells, or plasma did not produce appreciable levels of H_2_MCZ, if any. Alternatively, pig tissue extracts efficiently reduced MCZ into H_2_MCZ, as found in human volunteers (**Fig. 2A**). The identity of the metabolite produced was confirmed by LC-MS/MS as having the same MS/MS transitions and retention times as the *p-*H_2_MCZ made by NaBH_4_ treatment (Kloss *et al*., 2017) (**Fig. 2A**).

This reaction was found to preferentially use NADPH as a cofactor, and occurred in the S9 and cytosolic fractions, but not in microsomal fractions. The cytosolic reductase appears to be ubiquitous, as H_2_MCZ was detected after incubating MCZ with various pig organ extracts (liver, lungs, intestines, and spleen; **Fig. 2B**). In pig liver cytosol (PLC) at a MCZ concentration of 4.38 µM, the enzymatic reaction followed Michaelis-Menten kinetics with an apparent Km of 2.77 µM and a V_max_ of 7.46 ng/mL/min. After extensive characterization, this assay was applied throughout our investigation of MCZ reductase(s).

### Incubation of macozinone in human tissues

*Ex vivo* incubation experiments were also performed with MCZ incubated in S9 and cytosol fractions of post-mortem human liver, lung, and ileum tissue specimens. The results of H_2_MCZ production over 50 min are shown in **Fig. 2C**, and compared to the incubation results with pig tissues (**Fig. 2B left**). Interestingly, the H_2_MCZ yield was highest in human lung S9 and cytosolic fractions, while in pig tissues, H_2_MCZ was mostly produced in the liver and intestine. As in pig, no H_2_MCZ was produced upon incubation with human microsomal and mitochondrial fractions, confirming that the reducing enzyme is cytosolic.

### Enzyme candidate screening

Building on current knowledge of the chemistry of MCZ and its HMC metabolite, the main categories of enzymes that we examined successively were: i) carbonyl-reducing enzymes (CRE), ii) enzymes participating in redox cellular homeostasis, iii) enzymes implicated in nucleophilic aromatic substitution, and iv) α,β-unsaturated ketone reductases. The enzymes investigated and their prototypical inhibitors are listed in **Supplementary Table S1**.

#### i) Carbonyl reductases

As the MCZ carbonyl group is replaced in H_2_MCZ by an enolic hydroxyl group, we hypothesized that an initial reduction would take place, with subsequent tautomeric and mesomeric rearrangements. Several cytosolic NADPH-dependent enzymes could theoretically reduce MCZ, including the abundant cytosolic carbonyl reductase 1 (CBR1) from the short-chain dehydrogenases/reductases (SDR) superfamily, as well as several enzymes of the aldo-keto reductase (AKR) superfamily (Malatkova *et al*, 2014; Malatkova & Wsol, 2014). However, incubation with the respective active recombinant enzymes CBR1, as well as AKR 1A1, 1B1, 1B10, and 1C3, yielded no H_2_MCZ.

Using the above optimized conditions, we then incubated MCZ in PLC in the presence of several representative inhibitors of the major reduction pathways reported in **Supplementary Table S1**. Dicumarol appeared to be the most potent, followed by curcumin, with IC_50_ of 0.107 and 12.8 µM, respectively. Lineweaver-Burk plots (not shown) were indicative of non-competitive inhibition. Dicumarol is the archetypal inhibitor of the drug-detoxifying enzyme NAD(P)H:quinone oxidoreductase 1 (NQO1, also known as DT-diaphorase) (Asher *et al*, 2006), *a priori* an attractive candidate because of its cytosolic location, use of NADPH as cofactor, and MCZ’s quinone-like structure. The NQO1 hypothesis was further supported by the marked decrease in H_2_MCZ production on incubation with curcumin (Tsvetkov *et al*, 2005), quercetin (Karakurt, 2015), as well as epigallocatechin-gallate (EGCG) (937.5 µM) (Pandey *et al*, 2020) and Cibacron Blue (CB) (16.1µM) (Prochaska, 1988), all known NQO1 inhibitors (**Supplementary Table S1**). However, incubation of MCZ with purified recombinant human NQO1, whose reducing activity was ascertained with its prototypical substrate duroquinone, failed to produce detectable levels of H_2_MCZ.

#### ii) Enzymes participating in redox cellular homeostasis

The phenolic compounds curcumin (Fang *et al*, 2005), quercetin (Lu *et al*, 2006), and EGCG (Wang *et al*, 2008) have also been reported to inhibit the selenium-containing thioredoxin reductase I, TrxR1. The TrxR1 hypothesis was investigated by incubating MCZ with purified rat liver TrxR1 (with reducing activity confirmed using the model substrate 5,5-dithio-bis-(2-nitrobenzoic acid) (DTNB)), but this revealed no H_2_MCZ production. Moreover, specific gold-containing TrxR1 inhibitors aurothiomalate (ATM) (Omata *et al*, 2006) or auranofin (AF) (Gandin *et al*, 2010) did not affect H_2_MCZ production in PLC at 20µM or 7.38µM, respectively, thus excluding the involvement of TrxR1 in MCZ dearomatization.

The glutathione reductase (GR) pathway, inhibited by curcumin and quercetin (Guller *et al*, 2021), is also a key driver of redox cellular homeostasis, suggesting a possible implication in MCZ reduction. Also supporting this hypothesis, the formation of H_2_MCZ in PLC was inhibited by menadione (Duthie & Grant, 1989) and 2-AAPA (Seefeldt *et al*, 2009), two known inhibitors of glutathione reductase, with IC_50_ 15.5 µM and 30.6 µM, respectively. For redox cellular homeostasis, GR frequently functions in concert with other enzymes involved in glutathione metabolism (see below).

#### iii) Enzymes implicated in nucleophilic aromatic substitution

Meisenheimer complexes typically constitute reactive intermediates of nucleophilic aromatic (S_N_Ar) substitutions (Meisenheimer, 1902). This occurs during the conjugation of reduced glutathione (GSH) by glutathione-S-transferase (GST) (Hayes *et al*, 2005) with the archetypal electrophilic substrate 1-chloro-2,4-dinitrobenzene (CDNB), yielding the final non-HMC 1-GS-2,4-dinitrobenzene (Tierbach *et al*, 2020). Conversely, *in vitro* experiments using mammalian GST gave HMCs as the final dead-end glutathionyl conjugate of 1,3,5-trinitrobenzene (TNB), as this compound lacks a leaving group at the *ipso* carbon(Bico *et al*, 1994). Such HMC metabolites of TNB have also been reported *in vivo* e.g. in moth larvae (Clark & Sinclair, 1988) and octopus (Liou *et al*, 2000). In fungal species, the explosive 1,3,5-trinitrotoluene (TNT) is similarly metabolized into several HMC derivatives (Khilyas *et al*, 2013). The degradation of TNT into reduced HMC forms has also been reported in a *Mycobacterium* species(Vorbeck *et al*, 1994). The bacterial enzymatic pathway driving hydride addition to the aromatic ring of TNT is mediated by pentaerythritol tetranitrate reductase (PETNR) in *Enterobacter cloacae*(French *et al*, 1998) and by the xenobiotic reductase (XenB) in *Pseudomonas fluorescens I-C* (Pak *et al*, 2000). Remarkably and unusually, this is achieved by direct reduction (by enzyme-bound FMNH_2_) of the substrate aromatic ring to produce the HMC (French *et al*., 1998). By contrast, in humans, or more generally in mammals, isolable HMC products had never been reported *in vivo* for any xenobiotic metabolism before benzothiazinones (Kloss *et al*., 2017).

Given such considerations, we first verified that the gut microbiota was not responsible for a presystemic dearomatization of MCZ into H_2_MCZ, which, after intestinal absorption, would constitute the major circulating metabolite identified in rat and human plasma. However, *ex vivo* incubation of MCZ at 37°C with feces from omnivorous and vegetarian human donors yielded no detectable H_2_MCZ. Incubation with mouse caecum content did not yield H_2_MCZ either.

Glutathione-S-transferase (GST) was of particular interest since these phase II enzymes play a key role in the detoxification of various endogenous and exogenous hydrophobic electrophilic compounds (Chasseaud, 1979). Also, GSTs are inhibited by dicumarol (Mays & Benson, 1992) and Cibacron blue (Ismail *et al*, 2021), both found to markedly decrease H_2_MCZ production in our PLC assay. The three electron-withdrawing groups (NO_2_, CF_3_ and ketonyl) in MCZ (**Fig. 1A**) create a strong electrophilic spot at the position *p-* to NO_2_, which may be conceivably deactivated into a putative glutathionyl-MCZ (MCZ-SG) intermediate. Screening for the presence of the putative MCZ-SG conjugate by incubating MCZ with PLC extracts was therefore performed by Orbitrap LC-HRMS in positive and negative ionization mode, by monitoring the *m/z* signals at 763.80 corresponding to the molecular weight of the putative glutathionyl-MCZ (C_30_H_40_F_3_N_7_O_9_S_2_). However, in no case, was detectable glutathionyl-MCZ found in the PLC (data not shown).

#### iv) α,β-unsaturated ketone reductases

In addition to its ability to metabolize explosive nitro compounds, the bacterial NADPH-dependent reductase PETNR is remarkable for utilizing a series of α,β-unsaturated compounds (e. g. cyclic enone) as oxidizing substrates. These range from simple structures (e.g. cyclohexen-1-one) to complex α,β-unsaturated compounds (e.g. steroids) (Barna *et al*, 2001). Considering this intriguing interaction between bacterial PETNR and animal steroid hormones or their derivatives, we examined whether progesterone and 4-androstene-3,17-dione, two α,β-saturated steroid PETNR inhibitors (Barna *et al*., 2001), would interfere with a putative mammalian counterpart that might catalyze MCZ reduction. Both steroids, as well as the α,β-saturated hydrocortisone, were able to inhibit H_2_MCZ production in PLC, with IC_50_ values of **∼**50 µM (**Supplementary Table S1**). Prednisone, another PETNR substrate, also inhibited to some extent the formation of H_2_MCZ in PLC, presumably by enzymatic competition.

The concerted electron-withdrawing effects of NO_2_- and CF_3_-groups render the position *para*- to NO_2_ (β relative to the keto group) highly electrophilic (**Fig. 1A**), thus making the double bond adjacent to the carbonyl group a virtual α,β-unsaturated carbonyl substrate.

The enzymatic hydrogenation of endogenous or exogenous α,β-unsaturated carbonyl substrates (i.e. α,β-ketoalkene) can occur in mammals via a wide range of reactions(Huang *et al*., 2014). Enzymatic double-bond reduction of α,β-ketoalkene xenobiotics was repor ted for LY140091, a 2-benzylidine-3-ketocoumaran derivative (Lindstrom & Whitaker, 1984), for the aryl propionic NSAID agent CS-670 (Itoh *et al*, 2008), for the anticancer drug hydroxymethylacylfulvene (Yu *et al*, 2012), and, more recently, for the acetylcholinesterase inhibitor BYZX (Yu *et al*, 2013). For all these compounds, however, the double bond of the α,β-ketoalkene is not engaged in an aromatic cycle conjugation. The enzymatic double bond reduction of the CS-670 α,β-unsaturated ketone in rat liver cytosol was shown to be mediated by leukotriene B4 12-hydroxydeshydrogenase/15-ketoprostaglandinΔ13-reductase (LTB4 12-HD/PGR), later named prostaglandin reductase 1 (PTGR1)(Itoh *et al*., 2008). PTGR1, a highly inducible, cytosolic homodimeric enzyme, catalyses the NADPH-dependent reduction of the 13,14 double bond (i.e. an α,β-unsaturated carbonyl) of prostaglandin 15-keto-PGE2 (Wang *et al*, 2021). PTGR1 plays a key role in the activation of antitumor acylfulvenes derivatives (Yu *et al*., 2012) and may also mediate double bond reduction in the α,β-ketoalkene compound BYZX (Yu *et al*., 2013).

Importantly, some of the widely used NSAIDs, which primarily act as cyclooxygenase inhibitors, also inhibit PTGR1 (Clish *et al*, 2001; Mesa *et al*, 2015). We thus tested the ability of three common NSAIDs, diclofenac, indomethacin, and niflumic acid, to inhibit MCZ dearomatization in PLC. These experiments showed that diclofenac and niflumic acid inhibited MCZ reduction with virtually the same IC_50_ (16.2 and 16.3 µM, respectively), while indomethacin was slightly less potent (IC_50_ 23.6 µM). The inhibition of MCZ reduction by NSAIDs strongly suggested that PTGR1 was a likely candidate for this dearomatization reaction. This assumption was also supported by the strong inhibition demonstrated in our PLC incubation experiments by dicumarol, another inhibitor of α,β-ketoalkene reductases (Yu *et al*, 2013).

### PTGR1 and PTGR2 mediate macozinone dearomatization

Definitive demonstration of PTGR1 mediating MCZ dearomatization was provided by incubating MCZ with pure recombinant human PTGR1, which yielded a single metabolite, *p*-H_2_MCZ, with the same chromatographic retention time and *m/z* transition as the H_2_MCZ produced by PLC incubation. Human monomeric PTGR2 (also known as zinc-binding alcohol dehydrogenase domain-containing 1, ZADH1) shares 40% homology with PTGR1 (Mesa Solis, 2016). Like PTGR1, PTGR2 catalyses NADPH-dependent reduction of the α,β-unsaturated carbonyl of 15-keto-PGE2, a key step of terminal inactivation of prostaglandins (Wu *et al*, 2008), but unlike PTGR1, it cannot oxidize the hydroxyl group of LTB4.

Incubation of MCZ with pure recombinant human PTGR2 also produced *p-*H_2_MCZ that had the same chromatographic retention time and *m/z* transition as the H_2_MCZ produced in PLC incubation (**Fig. 2A**). Recombinant PTGR2 provided a three-fold higher yield of *p-*H_2_MCZ as compared to PTGR1 in the same incubation conditions.

Therefore, though not obviously apparent on considering their chemical structures, MCZ shares with prostaglandins an α,β-unsaturated carbonyl group (**Fig. 3A**), which likely explains why this xenobiotic is a substrate of enzymes catalyzing the reduction of 15-keto-PGE2 prostaglandins. Such reaction is unique since it implies enzymatic β-reduction of a double bond engaged in an aromatic ring, creating the unprecedented HMC. The proposed biochemical reaction of NADPH-dependent reduction of MCZ by PTGR1 and PTGR2 into H_2_MCZ is shown in **Fig. 3B**.

**Figure 3.**
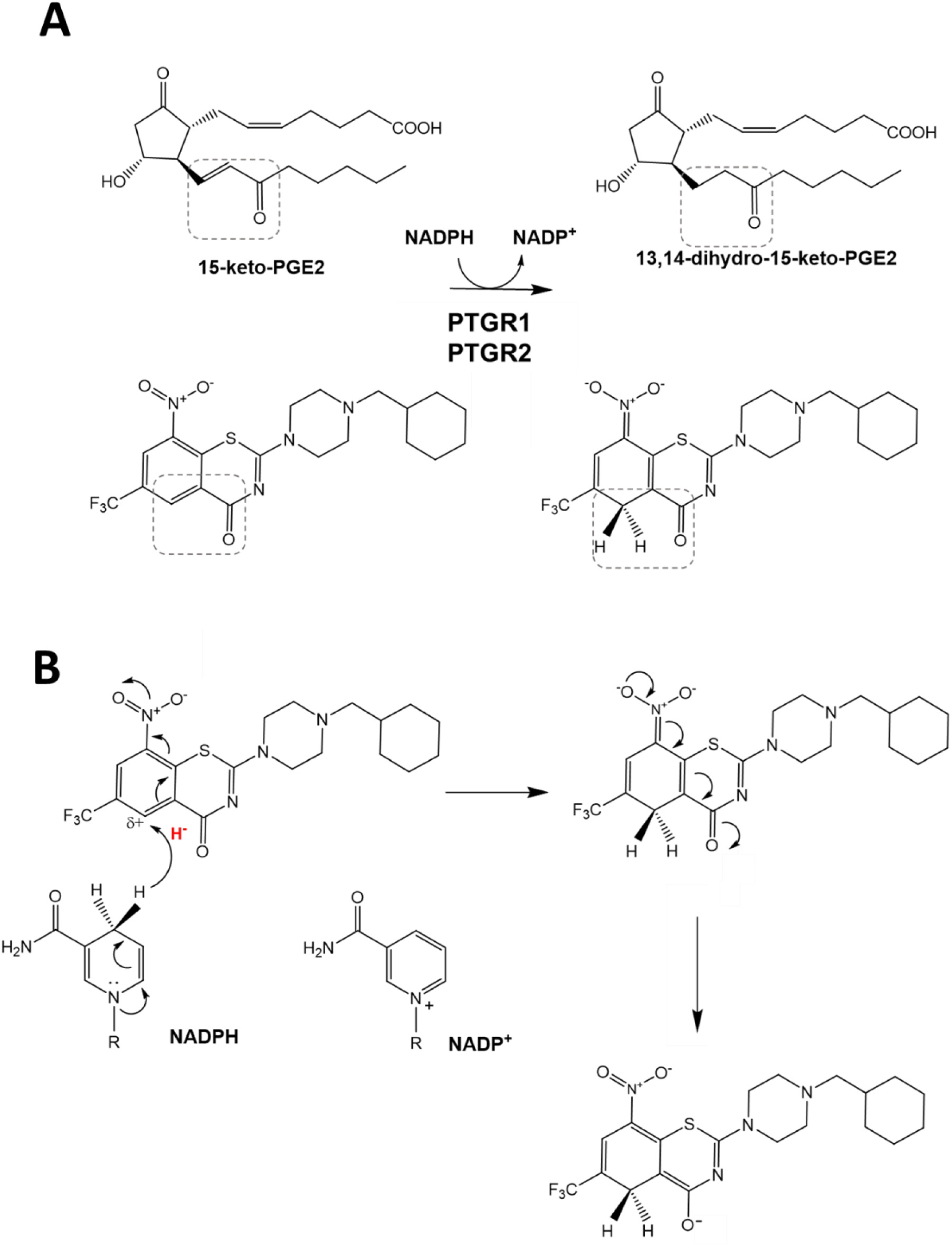
PTGR1 and PTGR2 are α,β-unsaturated ketone reductases that dearomatize MCZ into H_2_MCZ. (**A**) MCZ shares with prostaglandins PGE2 the α,β-unsaturated carbonyl moiety (dashed frame), owing to the presence of a strong electrophilic β carbon (H_2_MCZ shown in the keto form but can be also depicted as the enolate mesomere) (see **Fig. 1**). (**B**) Proposed mechanism of the NADPH-dependent reduction of MCZ by PTGR1 and PTGR2.

### Confirmatory *in vitro* enzymatic inhibition studies

Using the recombinant PTGR1 and PTGR2, we compared the relative potency of various representative inhibitors on enzymatic reduction activity. Concentration response profiles for PTGR1 and PTGR2 inhibition are shown in **Figs. 4A** and **4B**, respectively. Dicumarol was the most potent inhibitor against both PTGR1 and PTGR2 with IC_50_ values of 0.33 and 0.058 µM, respectively. Other NSAIDs such as diclofenac (Clish *et al*., 2001; Mesa *et al*., 2015; Mesa Solis, 2016) and indomethacin (Clish *et al*., 2001; Wu *et al*., 2008) also blocked PTGR1- and PTGR2-mediated MCZ reduction as did the specific PTGR2 inhibitor PTGR2-IN-1 (Parker *et al*, 2017).

**Figure 4.**
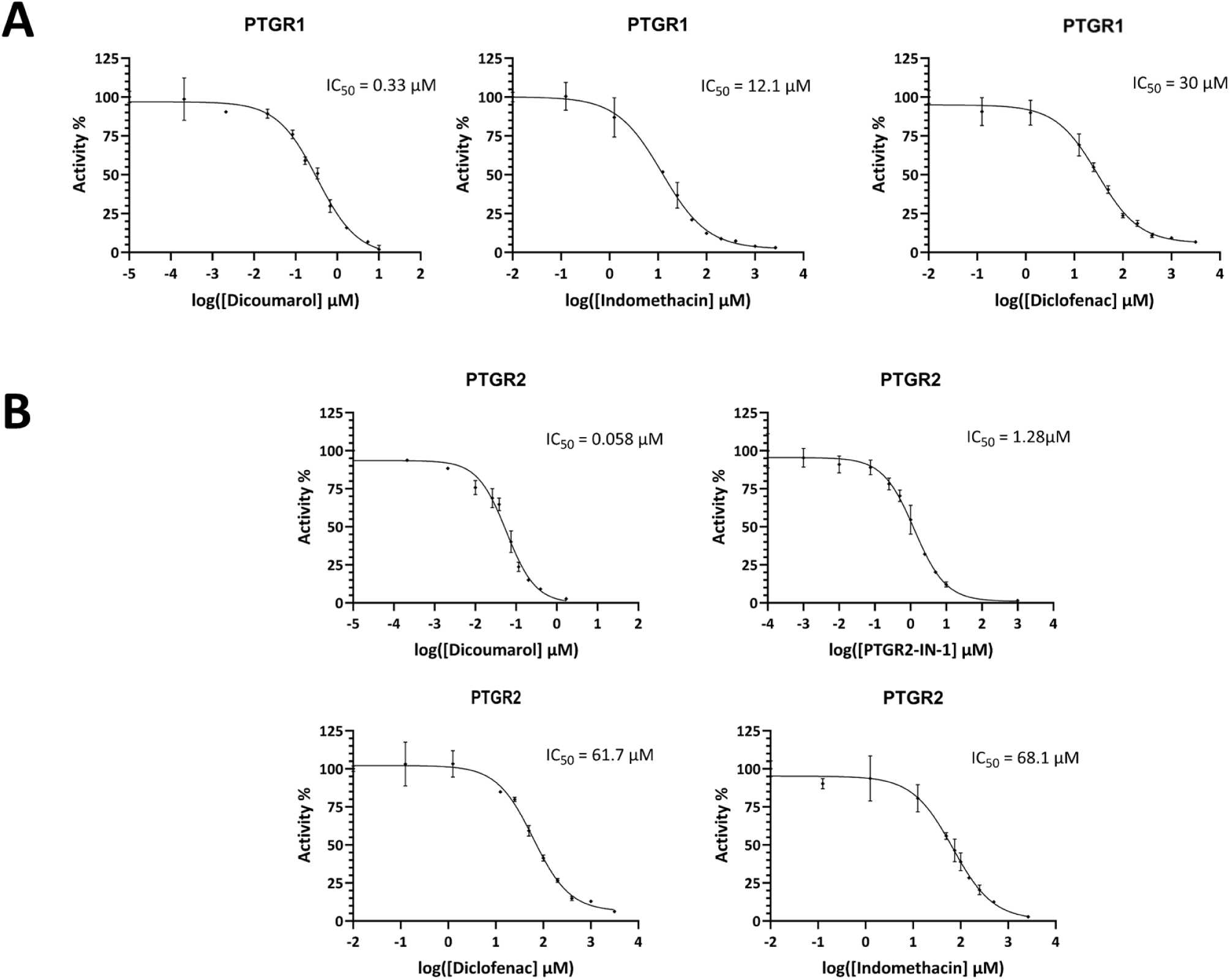
*In vitro* inhibition studies with human recombinant enzymes. Dearomatization mediated by (**A**) PTGR1 and (**B**) PTGR2, can be inhibited.

### Functional consequences of macozinone dearomatization

We then examined the antimycobacterial consequences of MCZ reduction through a combination of *in vitro* and *ex vivo* experiments, aiming to determine whether MCZ, its reduced metabolite or both are responsible for antimycobacterial activity. The controversy about the uncertain role played by H_2_MCZ in overall *in vivo* antimycobacterial activity was fueled notably by Liu et al(Liu *et al*., 2021), who argued that MCZ may actually be a prodrug for the active electrophilic moiety H_2_MCZ.

### *In vitro* MIC determination

*In vitro*, the synthetic mixture of *o*- and *p*-H_2_MCZ revealed a potency slightly lower than MCZ against two *Mycobacterium* species (**Supplementary Fig. 1A**), with an MIC of 1.2 µg/L against both *M. tuberculosis* H37Rv and *M. smegmatis* mc^2^155, compared with the MIC of 0.3 and respectively 0.6 µg/L known for MCZ in this assay.

Similarly, *in vitro* data with the BTZ-resistant MN84 strain (bearing a C394S mutation in DprE1) showed cross-resistance to MCZ and H_2_MCZ (**Supplementary Fig. 1A**). Also, in *M. tuberculosis*-infected macrophage experiments, MCZ and H_2_MCZ were of similar potency, indicating that H_2_MCZ enters human cells (or at least macrophages; **Supplementary Fig. 1C**).

However, in human sera samples spiked with known concentrations of MCZ, H_2_MCZ, or a mixture of both, the lowest MICs were found for serum samples spiked with MCZ followed by the MCZ/H_2_MCZ mixture, while the highest were for H_2_MCZ alone (**Fig. 5A**).

**Figure 5.**
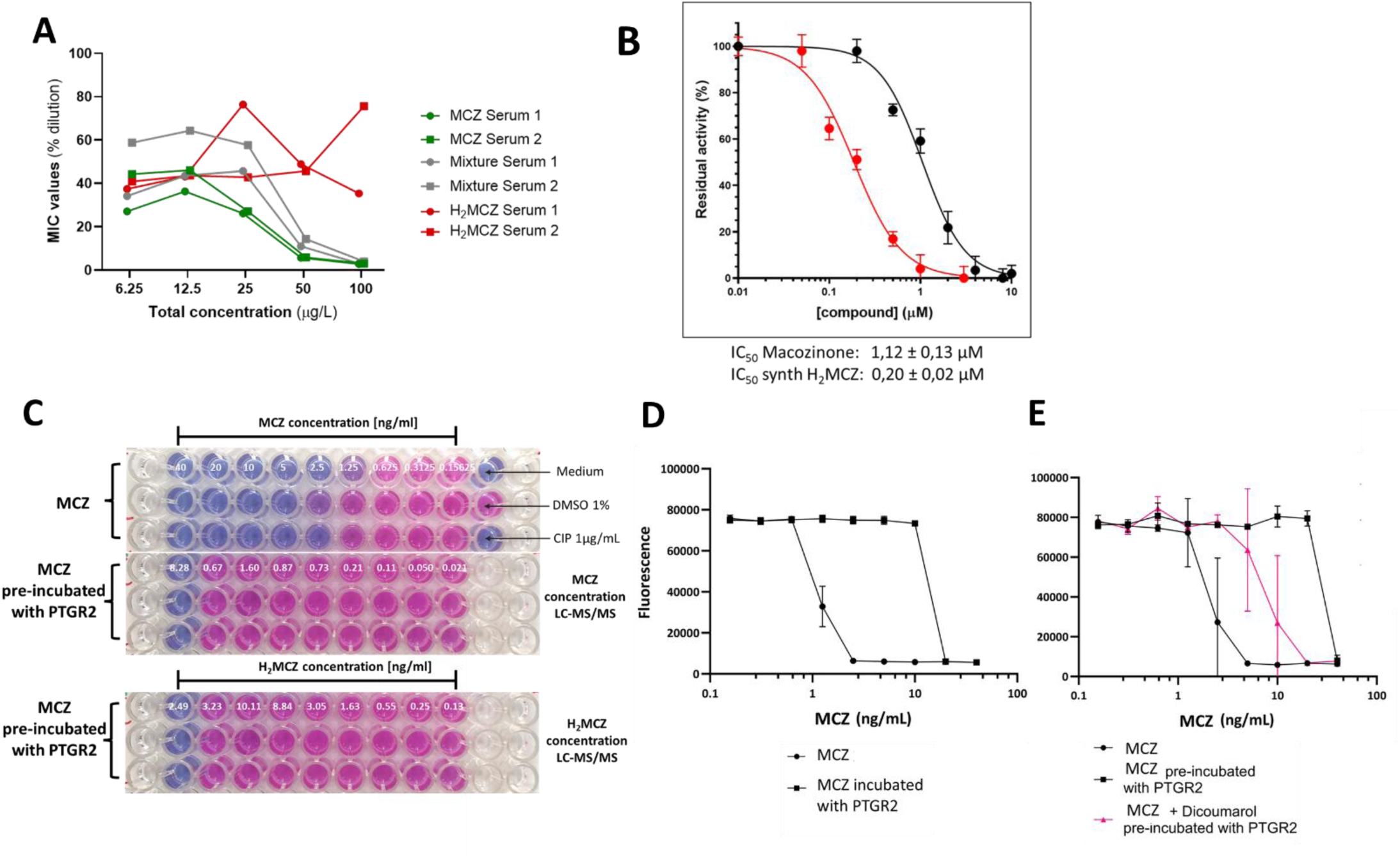
Functional consequences of macozinone dearomatization: *in vitro*. (**A**) Spike-in experiments. MIC values for human serum controls 1 and 2 spiked with MCZ (green), synthetic H_2_MCZ (red) or a mixture of the two (grey) at increasing concentrations. (**B**) In vitro DprE1 inhibition experiments. IC_50_ determination with pure enzyme. (**C**) Functional consequence of MCZ dearomatization. OD 0.0001; 72h-incubation, followed by 3h incubation with resazurin. PTGR2 concentrations: 2.4 µM. (**D**) Reduced MCZ antimycobacterial activity follows enzymatic dearomatization. MCZ and H_2_MCZ concentrations were determined by LC-MS/MS in the PTGR2-preincubated MCZ. (**E**) Decrease of macozinone antimycobacterial activity upon enzymatic dearomatization, mitigated by PTGR2 inhibition.

### Inhibition of DprE1

Then, we compared in a cell-free system the respective inhibition by MCZ and H_2_MCZ of the purified *M. tuberculosis* DprE1. Remarkably, in this *in vitro* test system, synthetic (*o-* and *p-*) H_2_MCZ was about 5-times more potent than MCZ itself (**Fig. 5B**).

### *In vitro* dearomatization of macozinone

The impact of MCZ reduction by PTGR2 on its killing activity against *M. smegmatis* is shown in **Figs Fig. 5C** (96-well plate, n=3) and **Fig. 5D** (concentration–activity profile). We used either MCZ (covering the relevant 10^-4^-10^2^ ng/mL concentration range) (**Fig. 5C**, upper three lines), or MCZ preincubated with PTGR2 (**Fig. 5C**, central and lower lines).

MCZ and H_2_MCZ concentrations were determined by LC-MS/MS in the PTGR2-preincubated MCZ wells, and are reported above the respective wells. PTGR2-mediated dearomatization of MCZ induced a clear shift of the blue-pink border (i.e. dead/alive bacteria) to the left, indicative of a decrease in antimycobacterial activity (**Fig. 5D**), corresponding to an increase in MIC from 1 µg/L with pure MCZ to *ca* 18 µg/L. This decrease in antimycobacterial activity after PTGR2-mediated dearomatization could be partially mitigated by adding dicumarol during the pre-incubation step (**Fig. 5E**).

### *Ex vivo* observations

We also examined *ex vivo* the antimycobacterial activity of plasma samples obtained from a representative volunteer participating in a 14-day multiple ascending dose phase 1 clinical trial with MCZ native crystal powder (https://clinicaltrials.gov/study/NCT03776500). Direct incubation of these plasma samples with *M. smegmatis* for 72h at 37°C was unsuccessful, as blank plasma *per se* already prevented growth. Instead, we carried out prior plasma protein precipitation of the volunteer samples, followed by dilution according to concentration quantified by LC-MS/MS, to bring MIC levels within the 20 - 0.02 ng/mL working range of our colorimetric assay.

**Fig. 6A** shows the PK profiles of MCZ and H_2_MCZ in the volunteer on the last day of the 14 day-multiple ascending dose regimen. **Fig. 6B** shows the corresponding MIC determination in the diluted extracts of plasma collected 0.5, 1, 1.5, 3, and 6h after oral MCZ intake, and **Fig. 6C** the respective antimycobacterial activity profiles, plotted against MCZ levels.

**Figure 6.**
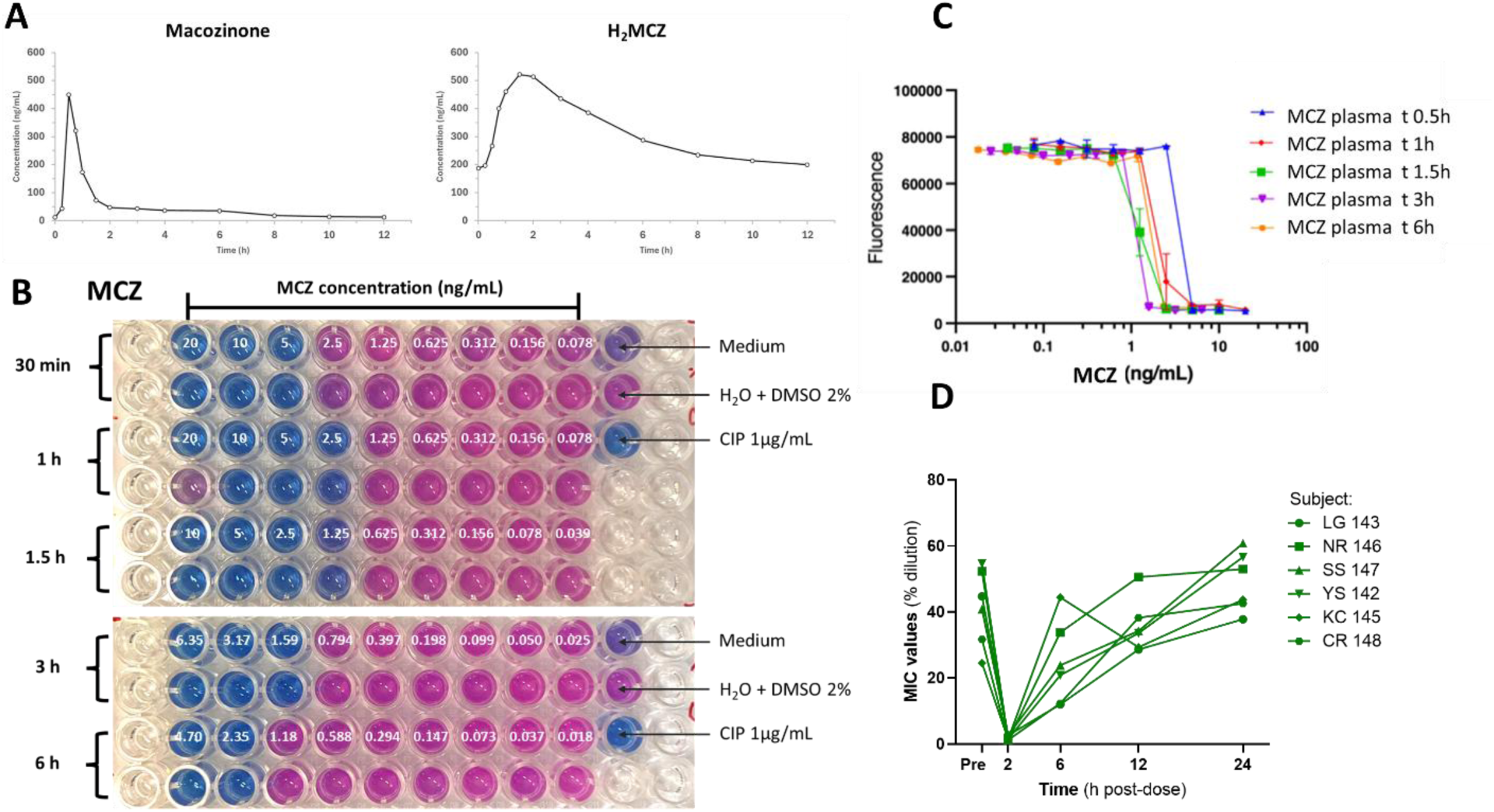
Functional consequences of macozinone dearomatization: *ex vivo*. (**A**) Pharmacokinetic profiles of macozinone and H_2_MCZ in plasma in one volunteer on the last day of the 14 day-multiple dose regimen. (**B**) Macozinone (MCZ) concentrations in wells. OD 0.0001 inoculum; 72h-incubation, followed by 3h incubation with resazurin. Serial dilutions of the volunteer’s samples on day-14 have been carried out according to plasma levels determined in the clinical pharmacokinetic study to bring the MCZ level in the working **∼** 0.02 µg/L - 20 µg/L range of the resazurin assay. (**C**) Antimycobacterial activity profile of plasma sample extracts from the same volunteer, plotted according to macozinone concentration. (**D**) MIC values for six subjects in the single ascending PBTZ169 dose study 1a, at various time points after dosing.

While the MIC follows the MCZ concentration closely, with the blue-pink border observed around 1 µg/L (**Figs. 6B** and **6C**), the H_2_MCZ concentration in the wells does not correlate with the antimycobacterial activity (**Supplementary Fig. S2**).

We also examined *ex vivo* the antimycobacterial activity of serum samples obtained from six volunteers included in a randomized, double-blind, placebo-controlled, single ascending dose trial of MCZ (https://clinicaltrials.gov/study/NCT03423030). **Fig. 6D** shows the MIC values of human serum collected at 0, 2, 6, 12 and 24h after receiving 320 mg of MCZ (Spray-Dried Dispersion formulation). All 6 subjects showed low (1-2 ng/mL) MIC values against *M. smegmatis* at 2h after dosing. For the pre-dose and all later serum samples, MIC values remained consistently high (35-70 µg/L).

## Discussion

Since the identification in 2017 of the metabolism of benzothiazinones into Hydride-Meisenheimer complexes (HMC) *in vivo*, the enzyme responsible for this reaction had remained mysterious. While HMC formation had never been reported in mammals, including humans, it has been widely described in microorganisms. This prompted us to direct our early investigations towards the gut microbiome as the source of H_2_MCZ production, but this hypothesis was incorrect.

Here, we report that the *in vivo* bioreductive dearomatization of MCZ into H_2_MCZ is catalyzed principally by the NADPH-dependent prostaglandin reductase 2 (PTGR2), and, to a lesser extent, by prostaglandin reductase 1 (PTGR1). PTGR1 and PTGR2 are key mammalian enzymes involved in the inactivation of the endogenous substrate prostaglandin (PG) 15-keto-PGE2. The multifunctional PTGR1 is responsible for the irreversible degradation of various eicosanoids, such as prostaglandins, leucotrienes, and lipoxins, endogenous lipid mediators crucial in a wide range of physiological and pathological processes, such as inflammation and defense responses, notably upon *M. tuberculosis* infection (Ning *et al*, 2023).

This is the first report of the implication of PTGR2 in the metabolism of a xenobiotic compound. The reaction is blocked by dicumarol, by the specific inhibitor PTGR2-IN-1, and by the widely used NSAIDs diclofenac and indomethacin. The related homodimeric PTGR1 also contributes to MCZ reduction and is inhibited as well by diclofenac, indomethacin, and niflumic acid. In fact, while all double bonds of the aromatic ring of MCZ are conjugated, the aromatic double bond adjacent to the carbonyl group is peculiar in that it can be considered as a virtual α,β-unsaturated carbonyl moiety, owing to the presence of a strong electrophilic β carbon (**Fig. 1A**). Therefore, though not obviously apparent on considering their chemical structures, MCZ shares with prostaglandins E the same α,β-unsaturated ketone chemical motif, though this is virtual for MCZ, as the double bond is part of the aromatic ring (**Fig. 2A**), which likely explains why this xenobiotic is a substrate of enzymes catalyzing the reduction of 15-keto-PGE2 prostaglandins. Such a reaction is unique in the sense that it implies enzymatic β-reduction of a double bond that is engaged in an aromatic ring, to create the HMC.

Conceptually, our findings expand the chemical repertoire of reactions catalyzed by mammalian reductases and uncover a noncanonical enzymatic mechanism by which host metabolism can neutralize antimicrobial agents. Notably, in contrast to the numerous cytochrome P450s and reductases generally involved in drug metabolism, it is most unusual for enzymes involved in the regulation of endogenous mediators, such as prostaglandins, to be also implicated in the metabolism of xenobiotics.

The discovery of PTGR1/PTGR2-mediated MCZ metabolism casts light on a critical metabolic reaction undergone by antimycobacterial benzothiazinones. Liu *et al* hypothesized that H_2_MCZ may be the active derivative of MCZ. In our hands, H_2_MCZ showed higher activity against purified mycobacterial DprE1 than the parent compound MCZ itself and in our *in vitro* MIC determinations, synthetic H_2_MCZ (*o*- and *p*-isomers mixture) was almost as potent as MCZ against *Mycobacterium* strains. However, we cannot exclude that the latter results may be compromised to some extent by spontaneous H_2_MCZ re-oxidizing to MCZ during the incubation process, in addition to the residual MCZ (evaluated at 7% by LC-MS/MS) present in the synthetic H_2_MCZ.

Indeed, LC-MS/MS analyses of the *Mycobacterium*-free incubation reaction medium reveal that after 72 h at 37°C, H_2_MCZ decreases by 93%, with a concomitant, yet not proportional, increase of MCZ (data not shown). Also, HMC species are commonly unstable intermediates, thus the electrophilic H_2_MCZ may react with some nucleophilic species present in the assay, which would explain the unbalanced stoichiometry of the H_2_MCZ oxidation reaction.

Overall, our data collectively indicate that the *in vivo* antimycobacterial activity is predominantly due to MCZ *per se*, rather than to its metabolite H_2_MCZ. This is supported by several preclinical experiments in mice that demonstrated strong anti-TB activity of MCZ. In mice, very low plasma levels of H_2_MCZ have been found (data not shown), in contrast to rats, dogs, and humans. Therefore, H_2_MCZ is unlikely to mediate any antimycobacterial activity in mice (Makarov *et al*., 2014).

Moreover, in our *in vitro* studies with MCZ pre-incubated with PTGRs before antimycobacterial tests, PTGR2-mediated reduction resulted in an overall decrease in activity (i.e. higher MIC). Likewise, the MIC values determined with serum samples spiked with MCZ, H_2_MCZ, or a mixture of both, or with *ex vivo* human plasma samples, follow the pharmacokinetic profile of MCZ, but not that of H_2_MCZ. Thus, we conclude that MCZ is not a prodrug and that H_2_MCZ, albeit circulating in abundance in humans, is not the active form. However, H_2_MCZ represents a drug reservoir thanks to its ability to reoxidize partly into MCZ, thus prolonging its pharmacokinetic half-life.

Exploiting their original whole cell assay, Kloss’s group has initiated optimization efforts to diminish the propensity of benzothiazinones to form HMC species, via the modulation of C-6 substituted nitro-benzothiazinone compounds, or by the replacement of the nitro-group, but this substantially decreased antimycobacterial potency. Instead, our work on the key role of PTGR1/PTGR2 for the dearomatization of benzothiazinones suggests that suitable enzymatic inhibitors of the reduction reaction (e.g. NSAIDs) could impair MCZ reduction and improve its exposure *in vivo*. Indeed, the addition of the inhibitor dicumarol to the macozinone-PTGR2 incubation mixture restored, in part, the antimycobacterial activity. Such an approach to boost exposure in patients is achieved by combining CYP3A4 inhibitors (ritonavir or cobicistat) with antiretrovirals or using the dehydropeptidase inhibitor cilastatin with the antibiotic imipenem. However, a word of caution is due about inhibiting PTGR activity, since endogenous eicosanoids are important modulators of the inflammation and immune response to *M. tuberculosis* infection. Notably, PTGR1 inactivates the inflammatory mediator leukotriene LTB4, which restricts inflammation and confers resistance to tuberculous infection (Tobin *et al*, 2013).

Despite the unequivocal demonstration of the PTGR1/2-mediated dearomatization of MCZ, several limitations in our work have to be acknowledged. First, only the *p-*H_2_MCZ isomer is observed upon incubation of MCZ with PTGR1/2 enzymes, as also seen in human plasma samples (**Fig. 2A**). Yet, in our *in vitro* tests, synthetic H_2_MCZ (i.e., *o*- and *p*-H_2_MCZ) was used, of which the naturally occurring *p*-H_2_MCZ constitutes only 41% (determined by ^1^H-NMR) of the isomeric mixture (**Supplementary Fig. 3**). Our efforts to separate the *o*- and *p*-H_2_MCZ by semi-preparative HPLC (using an adaptation of the method shown in **Fig. 2A**) were unsuccessful, as a substantial percentage was found to reoxidize to MCZ during the separation. In contrast, *p*-H_2_MCZ was found to be rather stable in volunteers’ plasma samples using appropriate precautions (Desfontaine *et al*., 2023).

Detailed pharmacokinetic aspects of the *in vivo* formation and elimination of H_2_MCZ remain to be described, ideally through a mass balance clinical trial. Proof of concept for the feasibility of boosting MCZ and BTZ-043 exposure by PTGR blockade is certainly warranted as it may allow use of lower doses of the drugs.

## Methods

### Chemicals, reagents, compounds, and other materials

Details of all reagents and products, including manufacturer information, are provided in **Supplementary information.**

### General

All incubation experiments with mammalian tissues have been performed at the Laboratory of Clinical Pharmacology of the University Hospital of Lausanne and the University of Lausanne, Switzerland. 1.5mL-Eppendorf tubes (Eppendorf Safe-Lock® Tubes were used for incubation experiments performed at 30°C and 37°C, using an Eppendorf ThermoMixer® equipped with the complementary heated lid Eppendorf ThermoTop, except for the incubation experiments with cellular fractions derived from the pig tissues, which were performed using a temperature-controlled water bath. Incubation at room temperature corresponds to 25°C.

Synthetic H_2_MCZ was prepared as follows: to a solution of 1.2 g MCZ (2.63 mmol) in 25 mL of acetonitrile with 0.5 mL of water, small batches of finely ground 0.2 g sodium borohydride (5.3 mmol) were added in small portions under argon with vigorous stirring at room temperature. A yellow-red precipitate forms after a few minutes. This solid was filtered and washed with a small amount of acetonitrile and diethyl ether and immediately vacuum dried. The yield is 0.8 g (66%), and the obtained product does not need recrystallization. The compound was stored in brown glass vials under argon at +4 °C. The synthetic mixture was determined to contain about 7-10% of residual MCZ by LC-MS/MS. The respective proportion of *o-* and *p-*H_2_MCZ in the synthetic mixture was determined by ^1^H-NMR (Supplementary **Fig. 3**) using the ratio of the aromatic proton (d 7.16 and 7.31 ppm, on C5 and C7, respectively) and was found by ^1^H-NMR (**Supplementary Fig. 3**) to contain 41% of naturally occurring *p*-H_2_MCZ and 59% of the *o*-isomer.

### Mass spectrometry methods

The LC-MS/MS analyses were performed using ultra-high-performance LC (UHPLC) instruments from ThermoFisher Scientific (San Jose, CA, USA) equipped with a binary pump, an autosampler with a flow-through needle, and a thermo-adjustable column compartment. A TSQ Quantis triple quadrupole MS was coupled to the UHPLC system via a diverter valve. A TSQ Altis triple quadrupole MS was coupled to the UHPLC system via a diverter valve. The ESI source was an Ion Max NG^TM^ electrospray ionization source equipped with a heated electrospray ionization (H-ESI) probe. HPLC vials containing sample extracts were stored at +4°C in the temperature-controlled autosampler over the entire analytical sequences. Extracts samples were kept in snap ring/crimp neck HPLC glass vial and compatible snap ring closures from Macherey-Nagel.

For enzymatic kinetic experiments, the concentration of H_2_MCZ produced was calculated using matrix-matched calibration curves containing H_2_MCZ at concentrations of 2, 5, 10, 20, 50, 100, 300, 800, and 2000 ng/mL. Matrix-matched calibrators samples contained a solution of NADPH generating mix (20 mM NADP⁺, 15 mM glucose-6-phosphate, 24 mM MgCl₂, in Tris-HCl 100 mM pH 7.4) and potassium phosphate buffer (incl. dithioerythritol, EDTA) (ratio 3:5, v/v), 5% DMSO:MeOH (1:1, v/v). Test samples and calibration were subjected to the same precipitation step with ACN (sample:ACN 3:10) prior to centrifugation (10min, 15’000g, +4°C), and insertion of supernatant (ca. 45µL) in HPLC vials. Of note, H_2_MCZ solutions are prone to degradation. It is therefore advised to prepare on the day of analysis stock and working solutions of H_2_MCZ to add *ex tempore* into blank plasma matrix for the preparation of calibration samples.

Details of other LC-MS/MS methods and Operating Procedures used throughout this work are available in the Supplementary Information.

### Tissue homogenization and subcellular fractions preparation

Pig organs, namely lung, liver, spleen, and intestine, have been obtained from a local butcher and immediately stored at -20°C after sacrifice, followed by long-term storage at −80°C. The organs were homogenized in an ice water bath in Tris-HCl 50mM pH 7.5 buffer containing 250mM sucrose, protease inhibitor mix (Thermo-Scientific), and EDTA (3mL of homogenizing solution per 1g tissue), using an Omni TH tissue homogenizer (Omni^®^).

A first centrifugation (10min, 600*g*, +4°C) of homogenized tissues allowed elimination of tissue debris pellets. The second centrifugation step (10min, 15’000*g*, +4°C) allowed the separation of mitochondria in the pellets, and supernatants constituting the S9 fractions. Ultracentrifugation (60min, 100’000*g*, +4°C) of the S9 fraction provides the cytosol (supernatant) and microsomes (pellets).

Mitochondria and microsome fractions were carefully washed using cold phosphate-buffered saline (PBS) to remove any remnant S9 or cytosol residues before storage at -80°C in homogenizing buffer and microsome storage buffer, respectively. Three to four careful washing steps were necessary to obtain colorless supernatants.

Protein concentration in fractions was determined using the bicinchoninic acid (BCA) assay (ThermoFisher Scientific, San Jose, CA-USA) using bovine serum albumin (BSA) as standard, with a working range of 25-2000µg/mL according to the manufacturer’s instructions. The high protein contents of some crude samples have necessitated preliminary tests with sample dilutions prior to the BCA assay.

### H_2_MCZ generation in tissues and cellular fractions

Tissue fractions, NADPH buffer, and potassium phosphate buffer (containing 0.1 mM EDTA and 5 nM dithioerythritol (DTT)) were pre-incubated for 10 min at room temperature (RT) to allow incubation mix equilibration. At T_0_, MCZ (final concentration 5.01µM) was added to the incubation mix and immediately transferred into an incubator at 37°C. At 5-, 10-, 20-, 35-, and 50-min time-points, a 15 µL incubation aliquot is collected and precipitated with 50µL cold acetonitrile (ACN) containing the stable isotopically labelled internal standards (ISTDs) MCZ_d11_ 80 ng/mL and H_2_MCZ-ISTD 40 ng/mL.

Precipitated samples are then centrifuged (10 min, 15’000*g*, +4°C) and the supernatant (*ca.* 45µL) is collected and analyzed by LC-MS/MS. A negative control sample containing no tissue is also tested in parallel. The final DMSO content in the incubation assay is 0.23%.

### H_2_MCZ production in pig cytosol or hemolyzed human blood in the presence of NAD(P)H

The cytosols from the four different pig tissues, namely liver, lung, spleen, and intestine, are diluted to reach a 2 mg/mL protein content in homogenization buffer. The pig cytosols, as well as non-diluted whole hemolyzed human blood (15 mL), are pre-incubated with either i) 6 mL NADPH 20 mM buffer, or ii) NADH 20 mM buffer, or iii) only Tris-HCl 100mM, in the presence of 10 mL potassium phosphate buffer (incl. EDTA, DTT) for 30 min at room temperature. The subsequent incubation procedure was the same as described above (final MCZ concentration 3.17 µM). A negative control sample (no cytosol) was also tested to ensure that H_2_MCZ was not formed spontaneously in the absence of cytosols. The final DMSO concentration in the assay was 0.12%. Incubation mixtures were extracted after 15 and 30 min and subjected to LC-MS/MS analyses.

### H_2_MCZ production in human cytosol and S9 fractions

Post-mortem human tissues were kindly provided by the NIH NeuroBioBank (Request #1628). Liver, lung, and ileum post-mortem specimens were obtained from a single 21-year-old donor (ID 695). The specimens were stored at –80°C. Sample preparation (tissue homogenization and S9 and cytosol isolation) was performed in our lab using the same procedure applied to pig tissues. Incubation experiments with MCZ followed the same procedure applied for pig samples as previously described.

### Pig liver cytosol enzymatic kinetics

The kinetics of the enzymatic reduction of MCZ with 20 mM NADPH were studied in PLC (adjusted to a protein concentration of 2 mg/mL), using the above-described incubation procedure and by varying MCZ concentrations from 0 µM to 30 µM. The maximum velocity V_max_ of the enzyme, as well as the apparent Michaelis constant K_M_, are determined from the Lineweaver-Burk plot of the reaction, calculated using Microsoft Office Excel 2016 and plotted with GraphPad Prism 9. Similar experiments were also performed by varying cytosolic pig liver protein concentration from 0 mg/mL to 20 mg/mL at a fixed 25 µM MCZ concentration in the medium (i.e. saturating).

### Enzymatic pathway screening assays

Macozinone dearomatisation inhibition experiments were performed by incubation of MCZ in PLC in the presence of prototypical inhibitors of various putative reductive enzymatic pathways (**Supplementary Table S1**). The experimental procedures used for studying the various enzymatic pathways are detailed in the Supplementary Information.

### Incubation studies with recombinant PTGR1 and PTGR2 enzymes

MCZ at a final concentration of 11.41 µM was incubated at 30°C for 15 min with 0.1 mg/mL recombinant human PTGR1 or PTGR2, in the presence of NADPH regenerating mix . Importantly, the incubation mixture also contained 6.25% EtOH to maintain MCZ solubility. Incubation mixtures were subjected to standard extraction with ACN solution containing ISTDs, and the supernatants were analysed by LC-MS/MS through monitoring the respective H_2_MCZ and MCZ *m/z* transitions. Both analyses were performed in duplicate.

### *In vitro* enzymatic inhibition studies

Recombinant human PTGR1 (3 µL, final concentration 1,000 nM, 36 µg/mL) or PTGR2 (2 µL, final concentration 250 nM, 10 µg/mL) were incubated with NADPH regenerating mix (8.5 µL for PTGR1 and 9.5 µL for PTGR2), β-lactoglobulin (4µL, final concentration 0.5 mg/mL) and inhibitors (dicumarol, indomethacin, diclofenac, or the PTGR2-specific inhibitor PTGR2-IN-1; 2 µL at different concentrations) for 10 min at room temperature (RT). As a control, the inhibitor was replaced by an equivalent volume of DMSO (inhibitor solvent). MCZ (2.5 µL, final concentration 12.5 µM) was then added and the incubation mixtures were incubated for 60 min at 37°C. To ensure MCZ solubility, the final incubation mixture contained 6.25% ethanol (v/v). After incubation, 15µL of the incubation mixture was precipitated on ice with 50 µL cold ACN containing isotopically labelled internal standards (MCZd_11_ 80 ng/mL and H_2_MCZ-ISTD 40 ng/mL). Samples were vortexed and then centrifuged (10 min, 17,130 g, 4°C), and the supernatant (40 µL) was diluted with Milli-Q water (2:1, v/v) and analyzed by LC-MS/MS, monitoring the respective H_2_MCZ and MCZ *m/z* transitions (the LC-MS/MS method is described in the Supplementary Information).

Enzymatic activity was normalized to the condition without inhibitor, which was defined as 100% activity, and relative activities were expressed as a percentage of this control. Half maximal inhibitory concentration (IC_50_) values were determined by non-linear regression (dose-response inhibition model) implemented in GraphPad Prism version 9 (GraphPad Software, San Diego, CA, USA), and error bars on graphs represent the standard deviation from the mean of technical duplicates.

### DprE1 inhibition assay

*M. tuberculosis* DprE1 was prepared as previously reported. Enzyme activity assays were performed at 30°C in a final volume of 100 μL, using a Amplex Red/peroxidase coupled assay slightly modified from Neres *et al(Neres et al., 2012).* The standard reaction mixture contained 50 mM glycylglycine at pH 8.5, 100 mM KCl, 0.2 μM horseradish peroxidase, 50 μM Amplex Red, 0.5 μM DprE1. The reaction was started by the addition of farnesyl-phosphoryl-*β*-_D_-ribofuranose (FPR) 400 μM, and the conversion of Amplex Red to resorufin was followed by measuring the increase in absorbance at 600 nm (ε = 54000 M^-1^ cm^-1^) on an Eppendorf BioSpectrometer.

For IC_50_ determination the enzyme activity assays were performed in the presence of different concentrations of inhibitors (0.05-10 μM), and values were determined by fitting data according to the following equation, using GraphPad Prism 8 software:

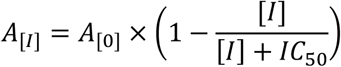

where A_[I]_ is the activity of the enzyme at inhibitor concentration [I] and A_[0]_ is the activity of the enzyme in the absence of inhibitor.

### *In vitro* dearomatization of MCZ

Recombinant human PTGR2 (2 µL, final concentration 2433 nM, 100 µg/mL) was incubated with NADPH regenerating mix (9.5 µL) and β-lactoglobulin (4 µL, final concentration 0.5 mg/mL), with or without inhibitors (dicumarol or 25 mM Tris-HCl buffer as negative control, 2 µL) for 10 min at RT. MCZ (2.5 µL, final concentrations ranging from 4.0 to 15.6 ng/mL) was then added and mixtures were incubated for 60 min at 37°C. To ensure MCZ solubility, the final incubation mixture contained 6.25% ethanol (v/v). After incubation, 65 µL of cold ACN containing isotopically labelled internal standards (MCZd_11_ 80 ng/mL, H_2_MCZ-ISTD 40 ng/mL) was added to the incubation mixture. Samples were vortexed thoroughly, and 64 µL was transferred to low-bind Eppendorf tubes. Samples were evaporated under nitrogen and reconstituted in 15 µL of dimethyl sulfoxide (DMSO), followed by 3 min sonication.

These pre-incubated samples were subsequently used for antibacterial activity testing, as described above, and the concentrations of MCZ and H_2_MCZ were determined by LC-MS/MS (described in the Supplementary Information).

### Antimycobacterial assays

The minimum inhibitory concentrations (MICs) of MCZ, H_2_MCZ, and rifampicin (RIF) were determined against *M. tuberculosis* H37Rv and *M. smegmatis* MC²155 using the resazurin microtiter assay (REMA) adapted from Palomino *et al*. (Palomino *et al*, 2002) The REMA assay was applied in the various experimental protocols described in the Supplementary information

### Ex vivo studies

The experiments shown in **Figs. 6A-6C** were carried out as follows. The plasma samples (collected on EDTA) were obtained from one volunteer included in a Phase Ib randomized, placebo-controlled, multiple ascending doses clinical trial (study name: IM-006–13, ClinicalTrials.gov identifier: NCT03776500) according to a protocol approved by the Institutional Ethics Committee. Briefly, 30 healthy volunteers received various doses of MCZ.HCl native crystal powder (NCP) ranging from 300 to 600 mg/day for 14 days, to assess the safety and tolerability of MCZ. Plasma (and urine) samples were collected at selected time points after MCZ intake on days 1 and 14 to study the pharmacokinetic/antimycobacterial profile of NCP-formulated MCZ in healthy volunteers.

Plasma samples from the volunteer on day 14 were analyzed to assess their antimicrobial activity against *M. smegmatis*. 100 µL-aliquot of plasma was precipitated with 300 µL of cold ACN. Samples were vortexed thoroughly and centrifuged (10 min, 17’130*g*, +4°C). A 300 μL supernatant aliquot was transferred to low-bind Eppendorf tubes. Samples were evaporated to dryness under nitrogen and reconstituted in Milli-Q water containing 8% (v/v) DMSO to obtain the desired concentration of MCZ and H_2_MCZ, followed by sonication for 3 min. These samples were used for the REMA assay. The REMA assay was performed as previously described, with the following modifications. For each plasma sample, 150 µL of inoculum (OD_600_ = 0.00013) was dispensed into the first wells of sterile flat-bottom 96-well plates, and 100 µL of inoculum (OD_600_ of 0.00010) was added to the following wells. 50 µL of each plasma sample was added to the first wells, mixed, and two-fold serial dilutions were performed across the plate. Each plate included a sterility control (7H9 broth only), a growth control (inoculum with Milli-Q water containing 8% DMSO), and a positive control (ciprofloxacin at 1 µg/mL). This experiment was performed in duplicate.

The *ex vivo* experiments shown in **Fig. 6D** were performed as follows. Briefly, serum samples were obtained from six volunteers included in a randomized, double-blind, placebo-controlled, single ascending dose to evaluate the safety, tolerability, pharmacokinetics, and *ex-vivo* antitubercular activity of PBTZ169 Spray-Dried Dispersion Formulation according to a protocol approved by the Institutional Ethics Committee. Details of experimental procedures for the assay are available in the Supplementary Information.

### Funding sources

This research was supported in part by the Bill and Melinda Gates Foundation (grant OPP1164837) through the study sponsor iM4TB (httaurops://im4tb.org/) and the Innovative Medicines Initiative 2 Joint Undertaking (JU) under grant agreement No 853989. The JU receives support from the European Union’s Horizon 2020 Research and Innovation Programme, EFPIA, Global Alliance for TB Drug Development Non-Profit Organisation, Bill & Melinda Gates Foundation, and University of Dundee (http://www.imi.europa.eu). Further support for this work and part of the salaries of DS, VD, FV, PT, and MB were also provided by a grant from the Swiss National Science Foundation (SNF) N° SNF 324730-165956 and SNF 324730_192449 to LAD. This analytical project carried out at the Laboratory of Clinical Pharmacology also benefits from the REQUIP grant SNF No 326000-121314/1 to LAD and from donations of the *Loterie Romande* for the acquisition of LC-MS/MS instruments.

## Acknowledgements

We are indebted to the NIH NeuroBioBank, for providing human post-mortem tissue specimens (Request #1628 to LAD)

We are grateful to the butcher ‘La Bouche-Rit, Chailly, Lausanne (Switzerland) (https://bouche-rit.ch) for providing pig tissues frozen immediately after animal slaughter.

Additionally, we thank M. Yves Laedermann and Ms. Catherine Brunner from Pharmacie Internationale Golaz in Lausanne, Switzerland, for providing some of the reagents, as well as Prof. Julijana Ivanisevic, Dr. Hector Gallart-Ayala, Ms. Rebecca Borreggine and M. Tony Teav from the Metabolomics Platform (University of Lausanne) for granting us access to their LC-HRMS equipment and providing insightful guidance.

We thank Gabriel Clerc, iM4TB (https://im4tb.org/) for invaluable operational work and fund management.

